# A new dual prokaryotic (*E. coli*) and mammalian expression system (pgMAXs)

**DOI:** 10.1101/2020.05.21.108126

**Authors:** Manabu Murakami, Agnieszka M. Murakami, Kazuyoshi Hirota, Shirou Itagaki

**Affiliations:** Department of Pharmacology, Hirosaki University Graduate School of Medicine, Hirosaki, 036-8562, Japan; Department of Anesthesiology, Hirosaki University Graduate School of Medicine, Hirosaki, 036-8562, Japan; Collaboration Center for Community and Industry, Sapporo Medical University, Sapporo, 060-8556, Japan

**Keywords:** protein, interaction, expression, plasmid

## Abstract

We introduce an efficient subcloning and expression plasmid system with two different modes (prokaryotic for expression in *Escherichia coli* with lac promoter and mammalian modes with cytomegalovirus promoter). The efficient subcloning (DNA insertion) is based upon a DNA topoisomerase II toxin-originated gene for effective selection with isopropyl-β-D-thiogalactoside (IPTG) induction. The new pgMAXs system is manageable size (4452 bp) and has also various types of protein tags (flag, myc, poly-histidine, Human influenza hemagglutinin, strep, and v5) for expression analysis. With pgMAXs system, various types of fluorescent proteins were subcloned and prtein expressions were confirmed. We also tried to identify epitope amino acid sequences for anti-calcium channel β2 antibody, by constructing epitope-library with DNaseI-partial digestion and subcloning into EcoRV site in pgMAXs. The new pgMAXs plasmid system enables highly efficient subcloning, simple expression in *E. coli* and that it has a simple deletion step of rare 8-nucleotide rare-cutter blunt-end enzymes for mammalian expression plasmid construction. Taken together, the pgMAXs system simplifies prokaryotic and mammalian gene expression analyses.

## Introduction

There are a number of commercial expression plasmids exist, for mammalian transient expression. The process of mammalian transient expression of a desired gene has mainly relied on a serial DNA recombination steps (two-step cDNA recombination): subcloning of the desired gene into a subcloning plasmid such as pBluescript (Agilent Technologies, Santa Clara, CA, USA), and subcloning the desired gene into a mammalian expression plasmid such as pcDNA3 (Thermo Fisher Scientific, Waltham, MA, USA) [1]. As this conventional method has conversion of a DNA fragment from one plasmid to another, we call it as C- (conversion) system. Each cloning step is often troublesome, due to the low efficiency of DNA ligation. In 2019, we established a novel pgMAX system, a dual expression system with two (prokaryotic and mammalian) expression modes [2]. This novel pgMAX system enabled efficient subcloning and gene expression in *Escherichia coli*(*E.Coli*) Furthermore, this system enabled simple and rapid construction of mammalian expression plasmid with its simple deletion-step (deletion of lac promoter unit with SwaI and PmeI). As this system needs only simple deletion of lac promoter unit and re-ligation, we name this type of plasmid system as D- (deletion) system.

The pgMAX system overwhelmingly simplified expression analysis, as it has practically only one subcloning step, while it has several disadvantages, such as relatively large plasmid size and only one tag-protein (flag) variety. Therefore, it has been desired to establish a new plasmid system with small size and different tag-proteins.

In the present study, we have established pgMAXs system, which has relatively small size (4452 bp) and various variants with different tag proteins (flag, myc, poly-histidine, Human influenza hemagglutinin, strep, and v5). The pgMAXs system simplifies prokaryotic and mammalian gene expression analyses.

## Methods

### Plasmid Construction

The novel pgMAXsflag was originated from former pgMAX [2]. DsRed2, pEGFP, pECFP and pEYFP plasmids were purchased from Clontech Laboratories (Palo Alto, CA, USA) [3]. For plasmid construction, PCR-based mutagenesis was performed. The conditions for PCR with high-fidelity Pfu DNA polymerase (Agilent Technologies, Santa Clara, CA, USA) were empirically modified (denaturation at 94 °C for 20 s, an annealing step at the calculated temperature (ca. 50 °C) for 30 s and an extension at 72 °C for 30 s, for 35 cycles). Amplified PCR products were gel-purified with a gel extraction kit (Macherey-Nagel GmbH, Dueren, Germany). The inhibitory unit (iUnit) was PCR-amplified from pgMAX with a specific oligo DNA (PmeIFor: gcggataacaatttcacagttt and XbaIiUnitRev: aaatctagacattcaggcctgacatttatat), digested with EcoRI and XbaI, and subcloned into EcoRI and XbaI sites in pgMAX, resulting in small plasmid size from 6125 bp to 4452 bp.

Various protein tag sequences (Myc, poly-Histidine, HA, v5 and strep) were introduced by PCR (MycFor: GATCCgaacaaaaactcatctcagaagaggatctgATg and MycRev: aattcATcagatcctcttctgagatgagtttttgttcG, HisFor: GATCCcatcatcatcatcatcatATg and HisRev: aattcATatgatgatgatgatgatgG, HAfor: GATCCTACCCATACGATGTTCCAGATTACGCTATg and HArev: aattcATAGCGTAATCTGGAACATCGTATGGGTAG, v5For: GATCCGgtaagcctatccctaaccctctcctcggtctcgattctacgATg and v5Rev: aattcATcgtagaatcgagaccgaggagagggttagggataggcttacCG, strepFor: GATCCagcgcttggagccacccgcagttcgaaaaaATg and strepRev: aattcATtttttcgaactgcgggtggctccaagcgctG).

Protocols using the pgMAX system as well as the entire pgMAX sequence have been deposited in protocols.io. (DOI dx.doi.org/10.17504/protocols.io.zq3f5yn). DNA ligation was done using standard ligation techniques (Takara DNA Ligation kit ver.2.1, Takara, Otsu, Japan). For the transformation, XL10-Gold ultracompetent cells (Tet^r^ Δ(mcrA)183 Δ(mcrCB-hsdSMR-mrr)173 endA1 supE44 thi-1 recA1 gyrA96 relA1 lac Hte [F’proAB lacI^q^ZΔM15 Tn10 (Tet^r^) Amy Cam^r^] (Agilent Technologies) were used.

A blunt-end DsRed2 DNA fragment was amplified using Pfu DNA polymerase with DsRed2-specific oligo DNA (DsRed2for: AaaGCTAGCatgGCCTC CTCCGAGAAC GTCATCA; DsRed2rev: aaaGAATTCagatctcaggaacaggtggtg). A blunt-end enhanced green fluorescent protein (EGFP) and its related ECFP and EYFP DNA fragments were amplified using high-fidelity Pfu with oligo DNA (ENFPfor: cccGCTAGCatgGTGAGCAAGGGCGAGGAG; ENFPrev: cccGGTACCGGCGGCGGTCACGAACTCCAG). The PCR-amplified product was inserted into the EcoRV site of pgMAXs.

### Cell Culture and Transfection

Cell culture and lipofection were performed as described previously [4]. Human embryonic kidney cells (HEK293, ATCC CRL 1573) were cultured in Dulbecco’s Modified Eagle’s Medium supplemented with 10 % fetal bovine serum. Exponentially growing cells were plated onto 35-mm dishes, and lipofection was performed using commercially prepared lipofectamine (Invitrogen, Carlsbad, CA, USA).

### Microscopy

Standard epifluorescence optics (Olympus, Tokyo, Japan) was used to visualize DsRed2 (excitation wavelength 563 nm, emission wavelength 582 nm) or EGFP (excitation wavelength 470 nm, emission wavelength 505 nm). DsRed2-related fluorescence was recorded with a Hamamatsu ORCA-FLASH 4.0 system (Hamamastu Photonics, Hamamatsu, Japan).

### Western Blot Analysis

For Western blot analysis using *E. coli*, 0.5 ml of the culture medium (LB medium at 37 °C, 12–16 h) was harvested and resuspended in 100 μl lysis buffer (1 mM EDTA, 1 mg/ml lysozyme) and incubated at room temperature for 15 min [5]. Aliquots (10 μl) of the homogenate from each clone were resolved by 15 % SDS-PAGE and subjected to Western blotting. A commercially available polyclonal antibody specific for the Ca^2+^ channel β2 subunit (Sigma-Aldrich, St. Louis, MO, USA) was used, followed by a secondary anti-rabbit IgG antibody conjugated to alkaline-phosphatase (Promega, Madison, WI, USA). The membranes were blocked in TBST (150 mM NaCl, 20 mM Tris–HCl [pH 7.5], 0.05% Tween-20) with 0.1% bovine serum albumin for 1 h, followed by incubation (16 h at 4 °C) in the presence of 10 pM TBST containing complete TM protease inhibitor cocktail (Roche Pharma, Basel, Switzerland). The membranes were washed three times with TBS (150 mM NaCl, 20 mM Tris–HCl [pH 7.5]) before AP activity was measured using the stabilized substrate for AP (Western Blue; Promega), as described previously [6].

### DNasel-partial deletion and expression analysis

To obtain randomly cleaved sequences from the C-terminal D-domain in the rabbit calcium channel β2a cDNA fragment (GenBank accession number X64297.1), D-domain sequence (717 bp) was PCR-amplified with specific primers (rB2Dfor: atggtagcagctgataaact and rB2Drev: gaattctcattggcggatgtaaacatc). The amplified DNA was partially digested with deoxyribonuclease I (DNase I), as previously discribed [7, 8]. The DNA fragments were blunted by klenow fragment (1 U) in the presence of dNTPs (0.1 mM each dCTP, dGTP, dTTP, 1 mM dATP) for 20 min at 37°C. DNA fragments were separated by electrophoresis (1.5 % agarose). DNA fragments were subcloned into 50 fmol of the pgMAXs, which had been cleaved with *EcoRV*. Colonies were plated on nitrocellulose filters (laid on LB plates containing 50 ug/ml ampicillin) at a density of 1~5 × 100 colonies per filter and incubated for 16 h at 37°C. Replicate filters were prepared and the filters were then blocked in 50 mM sodium phosphate, pH 7.4, 150 mM NaCl (PBS) containing 0.1% (v/v) Tween 20 and 5% bovine-serum albumin (BSA) and washed twice with the same solution. For screening with the commercially available anti-calcium channel β2 antibody (Sigma-Aldrich) the filters were incubated overnight at 4°C TBS (150 mM NaCl, 20 mM Tris–HCl [pH 7.5]) containing 0.1% (v/v) Tween 20 and 1% (w/v) BSA. The filters were washed three times with TBS (150 mM NaCl, 20 mM Tris–HCl [pH 7.5]) containing 0.1% (v/v) Tween 20, and further washed TBS (150 mM NaCl, 20 mM Tris–HCl [pH 7.5]) before AP activity was measured using the stabilized substrate for AP (Western Blue; Promega), as described previously [9]. The recombinant plasmids of positive clones were isolated by standard methods and cDNA inserts were sequenced.

### Statistics

Data are expressed as the means ± the standard error of the mean. Prior to statistical analyses, data were analyzed with the Shapiro-Wilk test. After confirmation of a normal distribution, statistical differences were further determined by Student’s *t*-test or an analysis of variance with a Bonferroni post hoc test. *P* < 0.05 was considered to indicate statistical significance.

## Results

### Plasmid Construction

Figure 1A shows the plasmid map of pgMAXs/flag (prokaryotic mode). The pgMAXs plasmid was based on pIRESpuro3 (Clontech). The pgMAX plasmid has two functional components, the prokaryotic and mammalian components. The prokaryotic component is for prokaryotic gene expression (lac promoter and lac operator) and for efficient subcloning with inhibitory unit (iUnit) (Fig. 1 prokaryotic unit) and inserted between CMV promoter and polyA tail sequence in the mammalian expression component. The iUnit originates from CcdB, a toxin targeting the essential DNA gyrase of *E. coli* [10]. This iUnit enables efficient subcloning, as plasmid with no insert will form no colonies. The prokaryotic component has SwaI and PmeI (both enzymes are 8 cutter and makes blunt-end) sites at its 5’-terminal and 3’-terminal ends, which enables simple deletion of prokaryotic component for constructing mammalian mode. At the PmeI site, a Kozak sequence followed by a Flag protein-coding sequence was inserted. A blunt-end DNA fragment can be inserted into the EcoRV blunt-end site within the multiple cloning site, which also contains an inhibitory unit (iUnit).

**Figure 1.**
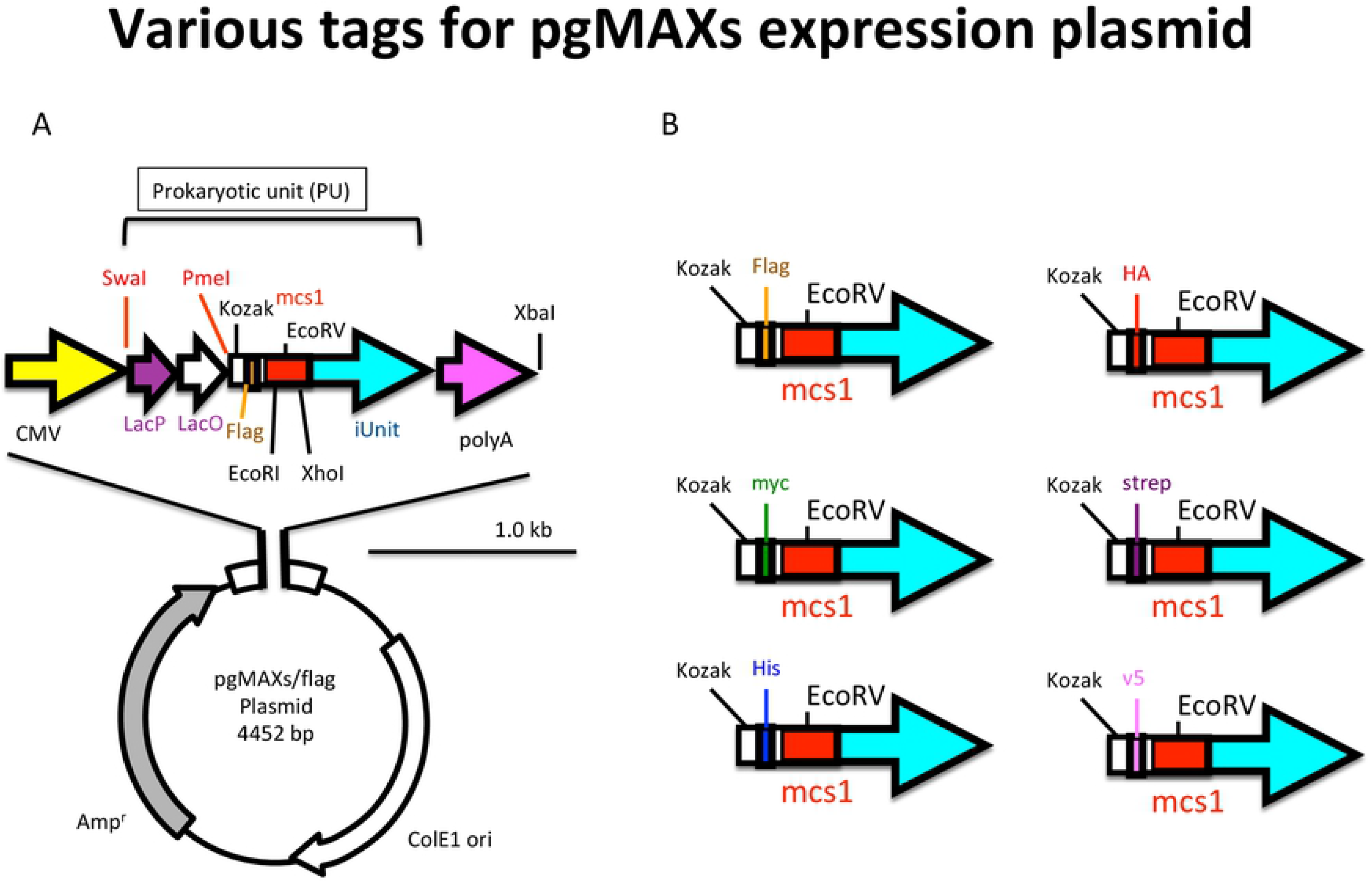
The pgMAXs plasmid system. **A. pgMAXs/flag construct** The pgMAX promotor has two functional components, prokaryotic and mammalian expression. The promoter is composed of a CMV promoter (yellow arrow) with a poly A tail (pink arrow)(mammalian unit). The restriction enzyme (SwaI, PmeI, EcoRI, EcoRV, XhoI and XbaI) sites are indicated. Oligo DNA for PCR screening is also indicated (pgMAXfor). **B. Various types of pgMAXs.** Construct with various tag proteins (myc, poly-histidine, HA, strep, and v5) from Kozak sequence until iUnit are indicated.

For protein expression analysis with different genes, it is often convenient to have different tag-proteins. Therefore, we constructed additional 5 different pgMAXs plasmids with myc, poly-histidine, Human influenza hemagglutinin (HA), strep, and v5 (Figure 1B). All constructs were successfully confirmed with IPTG-induced negative selection (no colony formation, data not shown).

### Simple Subcloning and Expression in Prokaryotic Mode

Various types of fluorescent protein (DsRed2, EGFP, ECFP, and EYFP) were PCR-amplified and inserted into the EcoRV site of pgMAXs/flag. For DsRed recombinant clones, after 16 h of incubation on LB agar plates containing ampicillin (150 μg/ml) and IPTG (1 mM for lac operon induction), colonies were observed under green light (excitation wavelength 563 nm) through a filter set (emission wavelength 582 nm). So far as DsRed recombination as concerned, among all of the 48 randomly picked colonies on the LB agar plates containing ampicillin and IPTG, 47 colonies contained the insert (DsRed) with an approximately 97.9 % success rate. Using fluorescence selection under green light (excitation wavelength 563 nm) through a filter set (emission wavelength 582 nm), all colonies (DsRed2: 5 of 5 colonies) contained the expected inserts with the desired sense-direction. Taken together, our data demonstrate that pgMAXs is a simple and universal cloning plasmid system for subcloning and prokaryotic (*E. colwas evaluated. For that purposei*) gene expression. We also analyzed colony numbers with former pgMAX (6125 bp) and pgMAXs/flag (4452 bp) for DsRed gene insertion. In four independent subcloning of DsRed fragment, pgMAXs formed more number of colonies (109.8 ± 16.2 and 31.8 ± 18.9* for pgMAXs and pgMAX, respectively; *\*P* < 0.05), which was expected due to its small size.

We also examined EGFP, ECFP and EYFP subcloning. For EGFP-related fluorescent proteins, blue light (excitation wavelength 470 nm) through a filter set (emission wavelength 505 nm) were used. Colonies with desired fluorescence were inoculated onto LB plates supplemented with ampicillin and IPTG for 16 h (Figure 2B). Each clones showed expected fluorescence, whereas pgMAXs/flag/ECFP tended to show lower ECFP expression, which might be related to the EGFP filter set to take picture.

**Figure 2.**
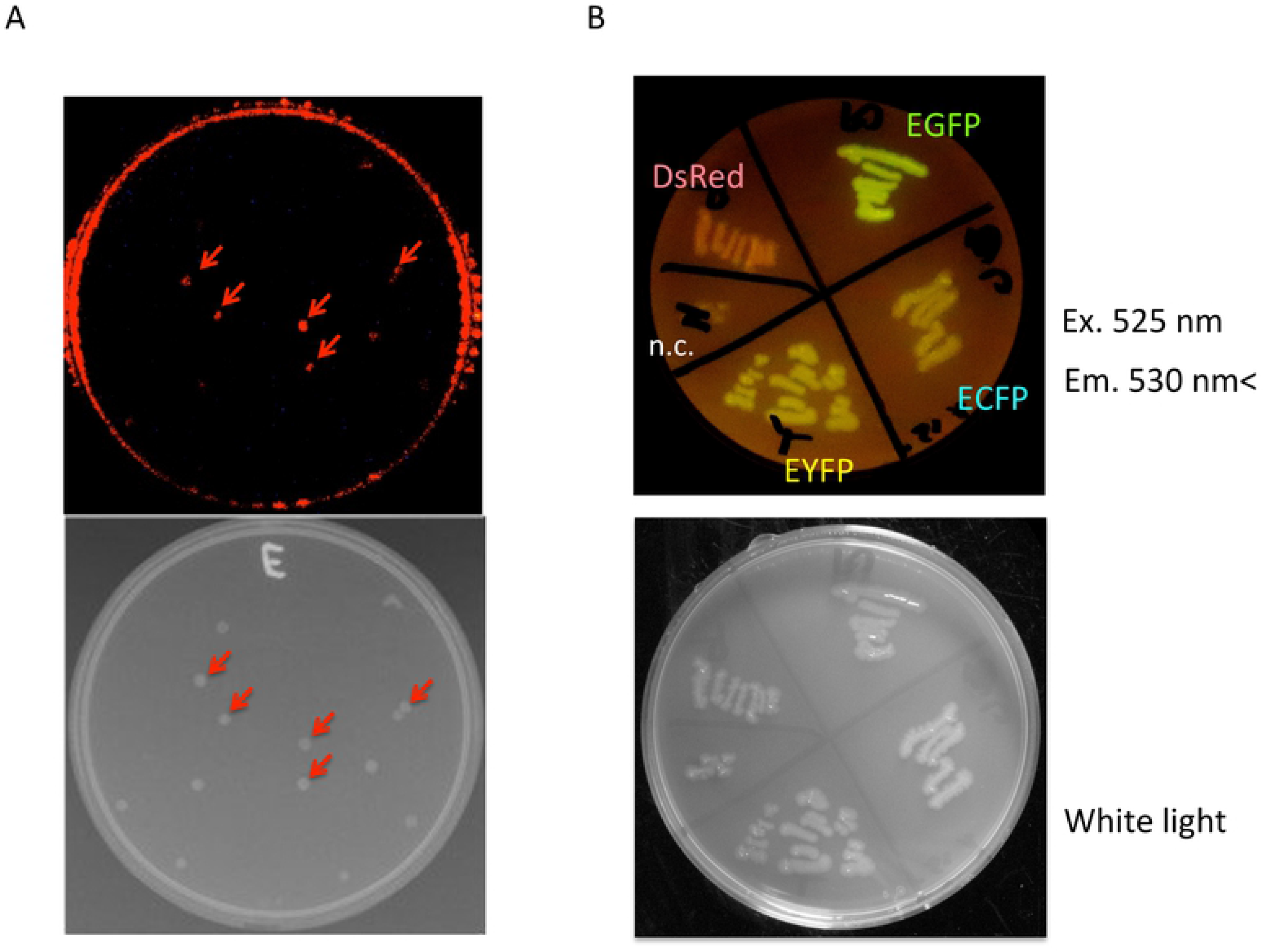
**A. Insertion of the DsRed2 fragment.** Recombinant clones of the pgMAX/blunt-end DsRed2 fragment were transformed into *E. coli* on LB plates supplemented with ampicillin and IPTG. Colonies observed under DsRed2 fluorescence under a green light and a red filter (upper panel) and a white light (lower panel) are shown. Five colonies with red fluorescence are indicated (red arrows). **B. Re-plated DsRed2, EGFP, ECFP and EYFP containing clones.** Fluorescent image under a blue light and a green filter, fluorescences of various proteins were observed (indicated, upper panel), while the negative control (pgMAXs without fragment) showed no fluorescence (n.c.). Image under a white light are shown (lower panel).

### Simple Preparation and Expression for Mammalian Mode

To analyze mammalian expression using the pgMAX system, the pgMAXs plasmid with the DsRed2 gene was further evaluated. Colonies with red fluorescence were selected and grown in LB medium supplemented with ampicillin (150 μg/ml) for 16 h, and plasmid DNA was purified using standard techniques. Insertion of the DsRed2 fragment was confirmed with restriction enzymes (EcoRI and XhoI). The purified plasmid DNAs of these red colonies were further restricted with SwaI and PmeI and re-ligated to delete the lac promoter unit (SwaI-lac promoter-lac operator-PmeI sequence), before being transformed with standard competent cells and cultured for 16 h at 37 °C. Plasmid DNA (mammalian mode) was further purified and used for transient expression in HEK293T cells.

After 24 h of plasmid DNA transfection in HEK293T cells, DsRed2-related fluorescence was readily observed with a fluorescence microscope (excitation wavelength 563 nm, emission wavelength 582 nm). The pgMAXs plasmids without DsRed2 fragments (negative control) did not exhibit DsRed2 fluorescence (5.0 ± 0.06 units [U], n = 8, Supporting Information Figure1A and B, pgMAXs, S1 Fig. 1), whereas red fluorescence was observed when the DsRed2 fragment was present (24.2 ± 0.8 U, n = 8, pgMAXs/Red). As a positive control, a DsRed2 or pgMAX/Red plasmids were also transfected, resulting in comparable fluorescence (18.5 ± 1.6 U, n = 8, DsRed2, and 18.0 ± 1.7 U, n = 8, pgMAX/DsRed,). The transfection efficiency of pgMAX/DsRed was comparable to that of DsRed2. The pgMAXs/DsRed transfected HEK293T cells showed a fair transfection rate (13.0 %; 19 of 146 cells, 48 h after transfection), compared to DsRed2 transfected HEK293T cells (11.3 %; 16 of 141 cells, 48 h after transfection). DsRed2-related fluorescence was further measured with the Orca system, confirming comparable fluorescence intensity with DsRed2 or pgMAX/Red plasmid transfection from day 1 to day 3 (S1 Fig. 1B), indicating useful application of novel pgMAXs system.

### Expression Analysis of an antibody recognition site

Furthermore, more generalized application of pgMAXs system for protein expression in *E.Coli* was evaluated. For that purpose, we tried to find epitope sequence of an antibody. Anti-calcium channel β2 antibody (Sigma-Aldrich) was used to recognize its epitope amino acid sequence. Full-length rabbit calcium channel β2a cDNA was divided into four (A, B, C and D) domains (Figure 3A) with four different oligo DNA pairs (S1 Table). Each domain was PCR-amplified and subcloned into the EcoRV site of pgMAXs. Each construct was readily transformed into *E. coli* and selected using ampicillin and IPTG. The expression of Flag-tagged sequences was analyzed with anti-Flag antibody (Figure 3Bi). The full-length β2a construct resulted in a major 75 kDa product, while additional small products were observed (Figure 3Bi, lane F). Other PCR amplified constructs formed expected protein bands (Figure 3Bi, lane A, B, C and D). Western blot analysis was performed using a commercially available polyclonal antibody specific for the Ca^2+^ channel β2 subunit (Figure 3Bii). Western analysis with anti-β2 antibody results in 75 kDa band in the lane F, which corresponds to full-length cDNA construct. In addition, anti-β2 antibody showed 31 kDa band in the lane D, which corresponds to the domain D (C-terminal domain), indicating that the domain D contains the epitope sequence for the antibody (Figure 3Bii lane D, arrow). Coomassie Brilliant Blue (CBB) staining of the total proteins on the gel was shown as control (Figure 3Biii).

**Figure 3.**
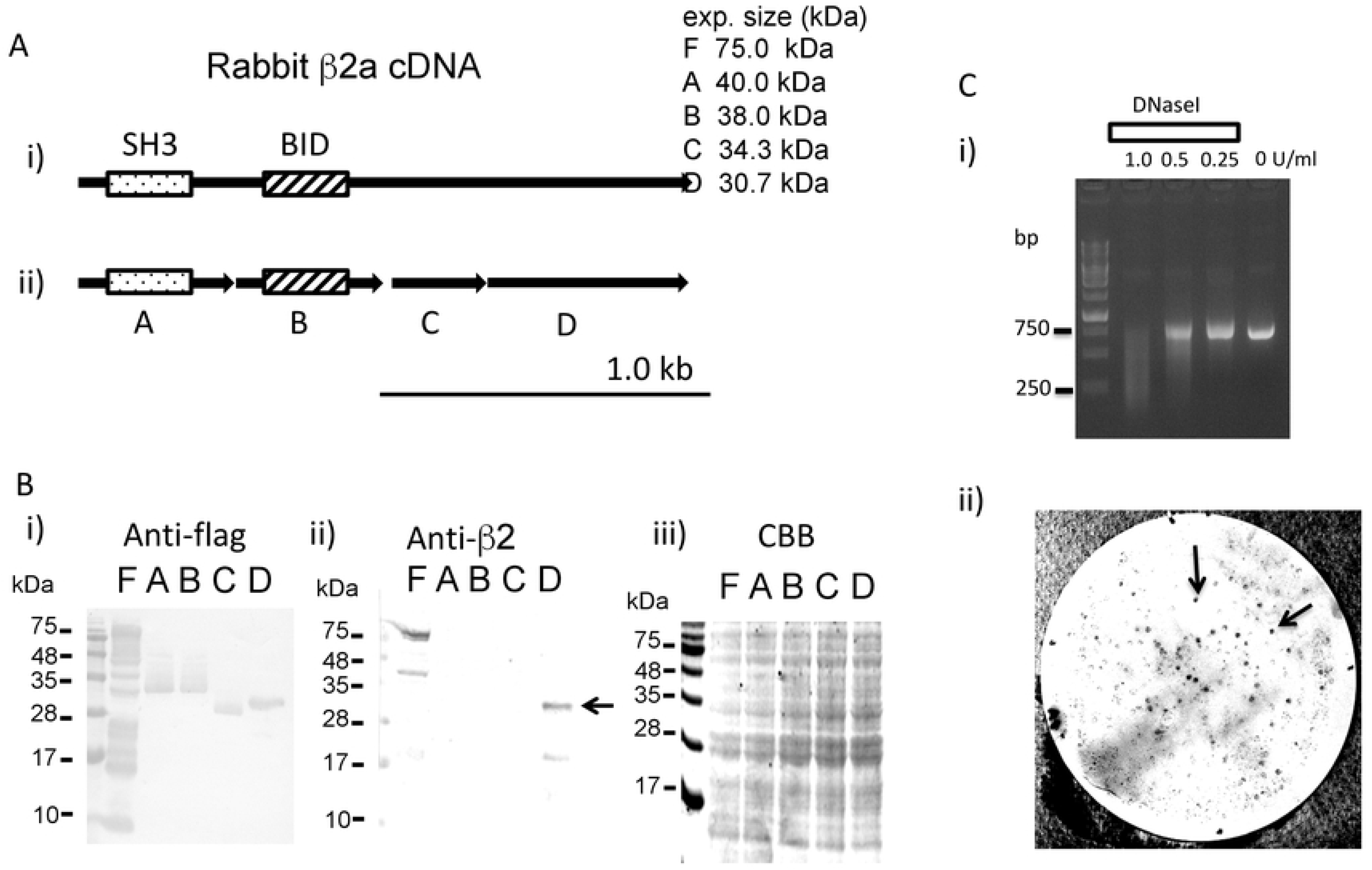
Expression analysis of a rabbit voltage-dependent calcium channel β2a subunit in *E. coli*. **A. Domain constructs of the rabbit voltage-dependent calcium channel β2a subunit.** Full-length cDNA clone of the rabbit voltage-dependent calcium channel β2a subunit (i) was divided into four domains (A, B, C and D; ii) with four different oligo DNA pairs (supplement Information Table 1). Scale bar = 1.0 kb. **B. Western analysis.** i) Immunodetection of the interactive domain of the anti-flag antibody. The names of the domains are indicated. The full-length clone showed different sized bands, while other constructs showed major single bands. The domains were indicated. ii) Immunodetection of the interactive domain of the anti-calcium channel β2 antibody. The full-length clone (F) and domain D contain the recognition site of the anti-voltage-dependent calcium channel β2 antibody (arrow). The names of the domains are indicated. ii) Coomassie Brilliant Blue (CBB) staining of the total proteins on the gel. **C. Randomly deleted epitope library screening.** i). DNA fragments electrophoresed on a 1.5 % agarose gel after treatment with 1,0.5 and 0.25 U DNase I per ml assay volume. An increase of the enzyme concentration leads to a decrease of cDNA fragment size. ii). Immunodetection of a nitrocellulose filter containing positive colonies isolated by screening a β2a subunit epitope library with the anti-calcium channel β2 antibody. Several positive colonies were observed (arrow).

### DNaseI-partial deletion of the domain D and expression analysis

D-domain sequence (717 bp) was PCR-amplified with specific primers (rB2Dfor: atggtagcagctgataaact and rB2Drev: gaattctcattggcggatgtaaacatc). The amplified DNA was partially digested with deoxyribonuclease I (DNase I), as previously discribed [7, 8]. The DNA fragments were blunted by klenow fragment (1 U) in the presence of dNTPs (0.1 mM each dCTP, dGTP, dTTP, 1 mM dATP) for 20 min at 37°C. DNA fragments were separated by electrophoresis (1.5 % agarose). DNA fragments were subcloned into the EcoRV site of the pgMAXs. Colonies were plated on nylon filters and replicate filters were prepared for immunoreaction. After the screening with the commercially available anti-calcium channel β2 antibody, we analyzed five independent positive clones (Fig. 3Cii). Sequence analysis revealed that all clones contained cDNAs in the appropriate reading frame and encoded peptide sequences derived from the D-domain of the ß2a subunit (Figure 4). The sizes of the peptide epitopes ranged from 108 to 144 amino acids (Fig. 4). In the Figure 4, overlapping amino acid sequence (441-480, dashed box) was indicated.

**Figure 4.**
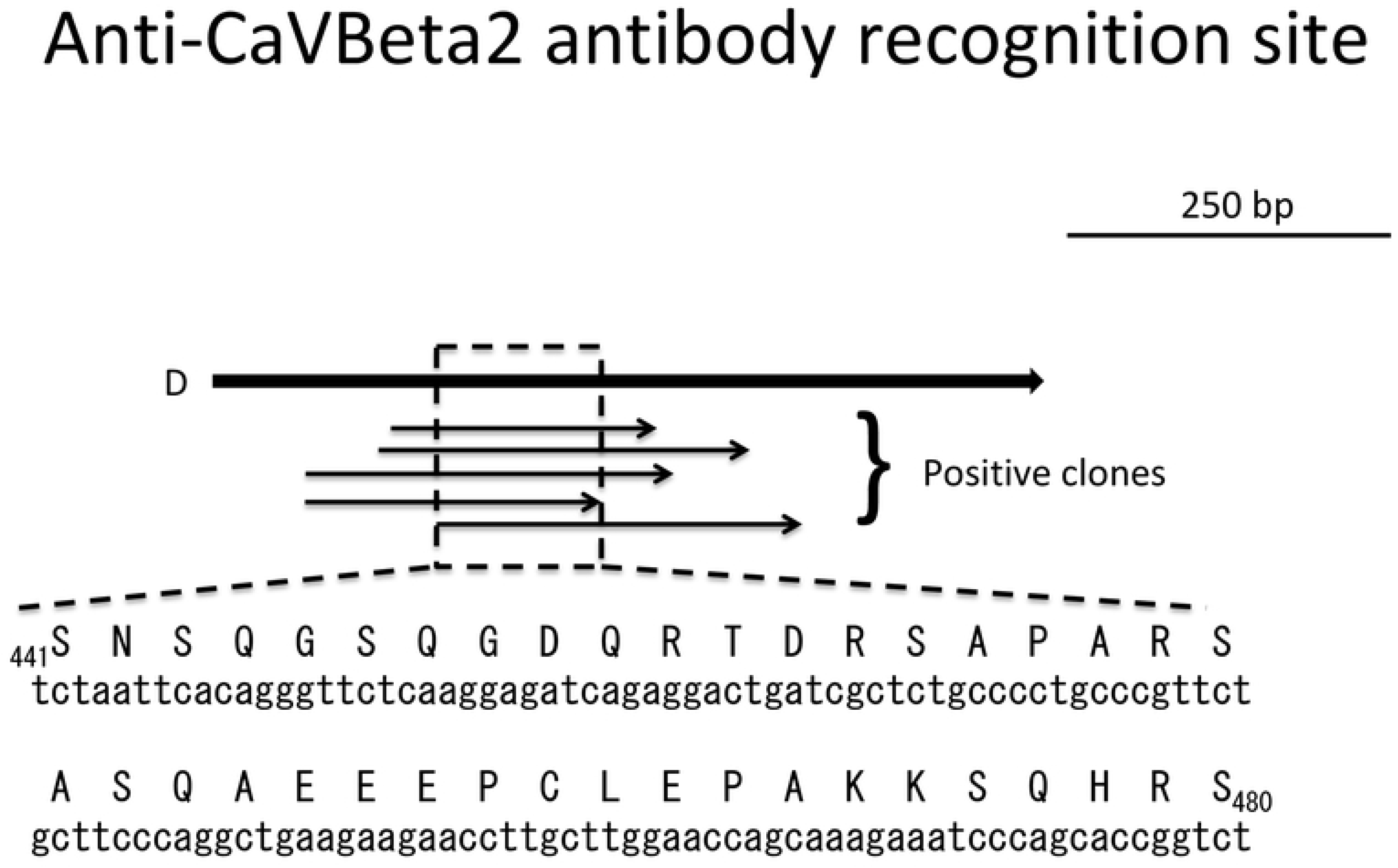
Identification of the anti-β2 subunit antibody binding site. Overlapping amino acid sequences from the five independent clones were indicated (dashed box). The first and last amino acids are numbered according to their location in the primary structure of β2a subunit.

Taken together, the pgMAXs plasmid resulted in highly efficient subcloning with prokaryotic expression and easy construction of mammalian expression plasmid vector with the desired gene expression.

## Discussion

In the present study, we established a new subcloning/expression plasmid (pgMAXs). This plasmid enables simple and highly efficient subcloning of a desired gene with standard techniques in *E. coli* (insertion step and prokaryotic mode), and easy construction of a mammalian expression plasmid within a few days (deletion step: restriction with SwaI and PmeI and re-ligation; after the deletion step, the plasmid is in mammalian mode). Recognition sequences of SwaI and PmeI are 8-nucleotide rare-cutter enzyme sites that are useful for achieving mammalian mode. In our analysis, the DsRed2-originating PCR fragment exhibited bright red fluorescence, indicating the establishment of a simple and efficient expression plasmid system. We further applied pgMAXs system for library construction for a protein-expression analysis to detect epitope sequence of an antibody and successfully identify epitope sequence.

After the establishment of former pgMAX, which contains IRES-puromycin resistant gene and has a relatively large size (6125 bp), several disadvantages has been recognized. For example, long pgMAX apparently shows a few numbers of colonies for blunt-ended DNA ligation. Because single cell experiment, such as patch-clamp analysis, is relatively limited application for a transient expression analysis, we deleted IRES-puromycin resistant gene with their vicinity sequences (1692 bp) from pgMAX, which results in novel pgMAXs (short; 4452 bp). Deletion of IRES-puromycin resistant gene resulted in three times colony formation (ca. 100 colonies instead of 30 colonies).

In addition, we prepared various tag proteins (flag, myc, poly-histidine, HA, strep, and v5) for protein expression analysis. With these various tag proteins, it could be possible to apply various types of expression analysis, such as protein-protein interaction.

Whereas pgMAXs enabled simple and easy conversion from prokaryotic to mammalian mode, it still needs DNA recombination to delete lac promoter unit, which locates between SwaI and PmeI. Previously, Udo has reported simple expression plasmid with modified lac expression sequences (lac promoter and operator), which do not need DNA deletion for prokaryotic and mammalian expression [11]. We also tried to establish expression plasmid with his sequences, but we could not get enough expression level of iUnit for DNA recombinant selection (unsuccessful selection between insert-containing clones and self-ligated clones, data not shown). Nevertheless, Udo’s concept (short and in-frame lac expression unit with desired gene) is interesting and should be examined in the future.

As expressed proteins in *E. coli* could be used for immunoblot analysis [7], we applied this plasmid system for protein analysis. Obviously, an example of high affinity interaction between two proteins is a binding between an antibody with its epitope sequence. Therefore, we tried to analyze epitope sequence of a commercially available anti-calcium channel β2 antibody. For that purpose, we first divided calcium channel β2 subunit into four domains (Figure 3). As C-terminal sequence (domain D, 717 bp) contains epitope sequence, we applied library construction with randomly deleted sequence of the domain D. With immunoreaction of the library with anti-calcium channel β2 antibody, we could identify epitope sequence, which indicates that the novel pgMAXs might be used for expression screening analysis for protein-protein inetraction.

In the present study, we used an iUnit originating from CcdB, a toxin targeting the essential DNA gyrase of *E. coli* [10]. As iUnit selection is quite effective for fragment ligation, pgMAX has significant superiority over classical DNA expression systems with its prokaryotic and mammalian expression modes, as the expressed protein can be examined after the fragment ligation. However, we think it is still possible to use other toxin sequences for this kind of selection, which should be analyzed in the future.

Since the discovery of restriction enzymes and DNA ligase, a number of plasmids have been established [12, 13]. Considering the use of restriction enzymes, the principal idea for the pgMAX plasmid might be not novel. Despite this similarity, as we mainly inserted 8-nucleotide rare-cutter enzyme sites (which should be cut every 4^8^ bp) at each end of a gene (lac promoter and operator, iUnit, IRES-puromycin resistance gene), each unit can be easily handled. Taken together, our results indicate that the pgMAXs plasmid system enables the simple and easy expression analysis of genes due to its efficient subcloning and rare-cutter sites.

## Conclusion

We established a fairly improved subcloning and expression plasmid system with two different modes (prokaryotic and mammalian modes) and various types of protein tags. The new pgMAXs plasmid system enables highly efficient subcloning of a blunt-end DNA fragment, simple expression in *E. coli* and that it has a simple deletion step for mammalian expression plasmid construction.

## Acknowledgements

This research was sponsored in part by Grants-in-Aid for Scientific Research from the Japan Society for the Promotion of Science, KAKENHI Nos. 17K08527, 17H04319 and 20K07255. No additional external funding was received for this study. We thank Mr. Maximilian Murakami for his technical advice.

## Author Contributions

Experiments were conceived and designed by MM. Experiments were performed by MM, AMM, KH and SI. Data analyses were performed by AMM and MM. Reagents, materials, and analysis tools were provided by SI and KH. The paper was written by MM, AMM, KH and SI.

## Additional Information

We declare no competing financial interests.

